# Phase separation of polyubiquitinated proteins in UBQLN2 condensates controls substrate fate

**DOI:** 10.1101/2024.03.15.585243

**Authors:** Isabella M. Valentino, Jeniffer G. Llivicota-Guaman, Thuy P. Dao, Erin O. Mulvey, Andrew M. Lehman, Sarasi K. K. Galagedera, Erica L. Mallon, Carlos A. Castañeda, Daniel A. Kraut

## Abstract

Ubiquitination is one of the most common post-translational modifications in eukaryotic cells. Depending on the architecture of polyubiquitin chains, substrate proteins can meet different cellular fates, but our understanding of how chain linkage controls protein fate remains limited. UBL-UBA shuttle proteins, such as UBQLN2, bind to ubiquitinated proteins and to the proteasome or other protein quality control machinery elements and play a role in substrate fate determination. Under physiological conditions, UBQLN2 forms biomolecular condensates through phase separation, a physicochemical phenomenon in which multivalent interactions drive the formation of a macromolecule-rich dense phase. Ubiquitin and polyubiquitin chains modulate UBQLN2’s phase separation in a linkage-dependent manner, suggesting a possible link to substrate fate determination, but polyubiquitinated substrates have not been examined directly. Using sedimentation assays and microscopy we show that polyubiquitinated substrates induce UBQLN2 phase separation and incorporate into the resulting condensates. This substrate effect is strongest with K63-linked substrates, intermediate with mixed-linkage substrates, and weakest with K48-linked substrates. Proteasomes can be recruited to these condensates, but proteasome activity towards K63-linked and mixed linkage substrates is inhibited in condensates. Substrates are also protected from deubiquitinases by UBQLN2-induced phase separation. Our results suggest that phase separation could regulate the fate of ubiquitinated substrates in a chain-linkage dependent manner, thus serving as an interpreter of the ubiquitin code.

**Significance:** Covalent attachment of polyubiquitin chains to eukaryotic proteins is a common protein quality control signal. Ubiquitination often marks proteins for degradation by the proteasome, but can also drive non-degradative outcomes. Proteins, including UBQLN2, that bind both polyubiquitin and the proteasome can either enhance or inhibit degradation. The ALS-related UBQLN2 is recruited to membraneless organelles, including stress granules, and undergoes phase separation *in vitro*, but the effects of phase separation on substrate fate are unknown. Herein we show that UBQLN2 phase separation is modulated by polyubiquitinated substrates in a linkage-dependent fashion. We show that two functional outcomes, degradation and deubiquitination, are differentially affected by phase separation. Our results suggest that phase separation of substrates and UBQLN2 could control protein fates.

## Introduction

Eukaryotic proteins can be post-translationally modified in many ways, with ubiquitination being one of the most common modifications (1). Ubiquitin (Ub) is typically attached to the amino group of a lysine on a substrate via an isopeptide bond, with additional Ubs attached to one or more lysines of the first Ub to form a polyubiquitin (polyUb) chain. The seven lysines and the N-terminal amine within Ub give rise to diverse polyUb chain architectures, which can lead to different fates within the cell (2-4). For example, K48-linked chains and some mixed linkage chains (K11/K48 and K48/K63) target proteins to the proteasome for degradation, whereas K63-linked chains are involved in DNA damage repair, and M1-linked chains are involved in immune response. These different polyUb chains make up the Ub code. However, our understanding of how these different chain types signal for different protein fates remains limited, especially since chain types with different *in vivo* fates (i.e. K48 and K63) can both be used to signal proteasomal degradation *in vitro*, with similar rates of degradation observed (5, 6).

An additional layer of regulatory complexity comes from a family of UBL-UBA shuttle proteins, which bind to the proteasome or other protein quality control machinery elements (via the UBL domain) and to ubiquitinated proteins (via one or more UBA domains) (7). In some cases, shuttle proteins facilitate protein degradation, but in other cases, they seem to be protective and actually prevent degradation (8-12).

We and others have recently shown that some of these shuttle proteins such as Rad23B, UBQLN2, and its homologs UBQLN1, UBQLN4 and yeast Dsk2 can undergo phase separation, a phenomenon in which proteins self-assemble via multiple interactions to form biomolecular condensates (membrane-free compartments) in the cell or liquid droplets *in vitro* (13-18). UBQLN2 forms stress-induced condensates in cells and can be recruited to stress granules, a type of stress-induced biomolecular condensate hypothesized to form via phase separation of RNA-binding proteins and RNA (15, 16). UBQLN2 mutations linked to frontotemporal dementia and amyotrophic lateral sclerosis affect phase separation behavior of UBQLN2 both in cells and *in vitro* (19-22). Importantly, Ub also affects UBQLN2’s propensity to phase separate, such that monoUb drives disassembly of UBQLN2 droplets *in vitro* in a concentration-dependent manner (16). Recent experiments with multiple polyUb linkage types revealed that length of the chain, linkage type, Ub-Ub spacing, and concentration are all important variables that modulate UBQLN2 phase separation (23, 24). Generally, K48-linked polyUb chains inhibit phase separation while K63-linked polyUb chains promote phase separation. In these situations, the valency of the polyUb chain and the arrangement of the Ub units can enable it to act as a scaffold and bring multiple shuttle proteins together to promote phase separation.

Although experiments with unanchored polyUb chains have revealed some of the rules that control phase separation of UBQLN2, we do not yet know how polyubiquitinated substrates behave in the presence of UBQLN2, or how substrate localization to phase separated condensates affects substrate fate (25). Using model substrates in an *in vitro* reconstituted system, we show that UBQLN2 and ubiquitinated substrates can reciprocally induce one another’s phase separation and that phase separation can protect substrates from both proteasomal degradation and deubiquitination.

## Results

To determine how substrate ubiquitination affects UBQLN2 phase separation, we ubiquitinated a model substrate consisting of R-Neh2Dual, an artificial degron derived from the N-terminus of Nrf2, followed by superfolder GFP (26) (sGFP; **Figure 1A**). The R-Neh2Dual degron can be ubiquitinated with a Ubr1 E3 ligase system, generating K48-linked chains; an Rsp5 system, generating K63-linked chains; or a Cul3/Rbx1/Keap1 system, generating branched ubiquitin chains containing K48, K63, and other linkages (27-30). We hereafter refer to these substrates as K63-linked, K48-linked or mixed.

**Figure 1.**
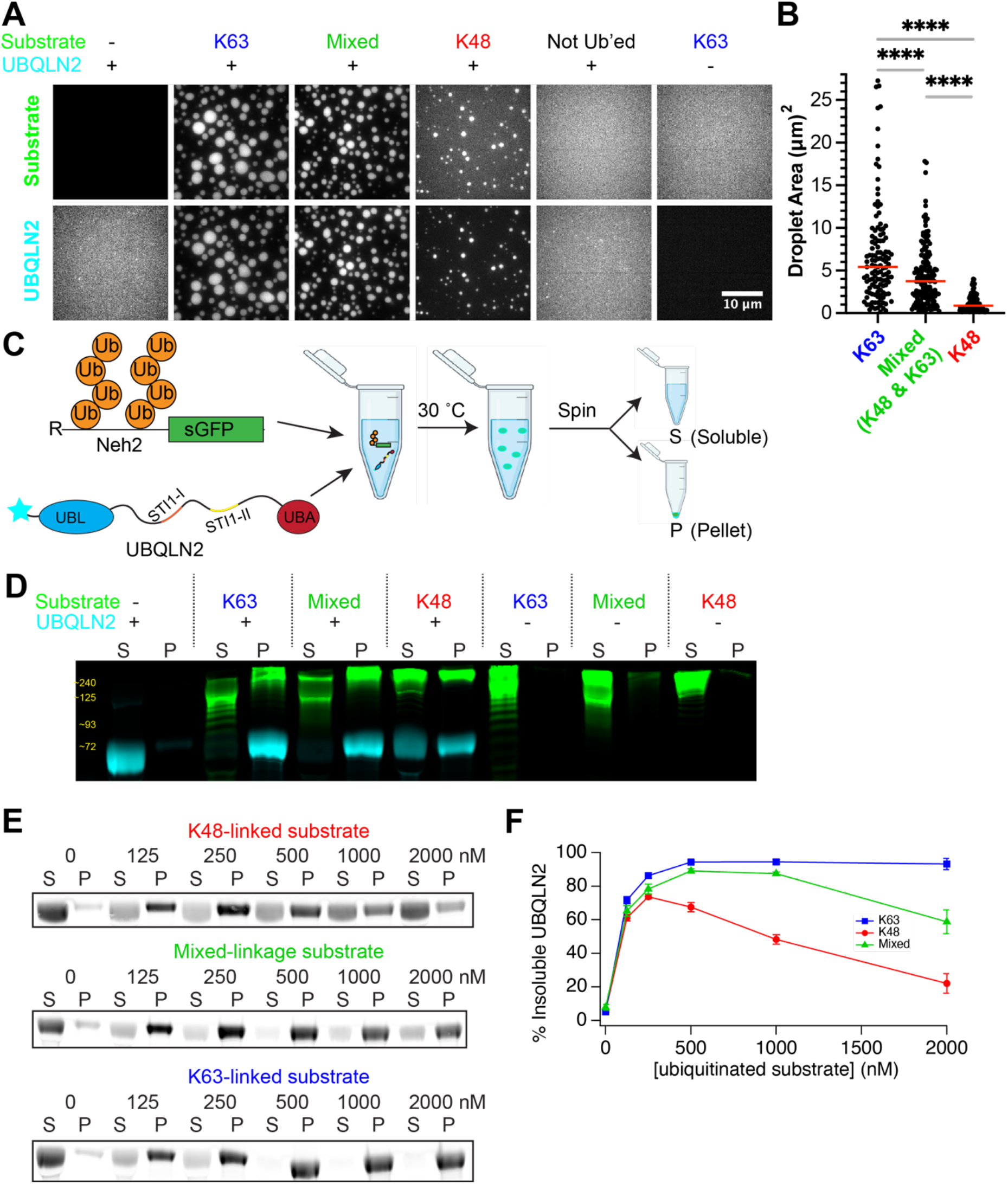
Ubiquitinated substrates phase separate and sediment UBQLN2 in a linkage-dependent manner. A) Fluorescence microscopy of 10 μM UBQLN2 (1% labeled with Alexa Fluor 647) incubated with or without 500 nM ubiquitinated (or not Ub’ed) substrate for 1 hr at 30 ºC. B) Quantification of droplet size from images in E (K63: n=125, Mixed: 161, K48: 102 droplets) and statistics applied with Welch’s two-tailed t-test (^****^, p < 0.0001). C) Sedimentation assays use R-Neh2Dual-sGFP substrate ubiquitinated with K63-linked (Rsp5), K48-linked (Ubr1), or mixed linkage chains (Keap1/Cul3/Rbx1). After mixing substrate with UBQLN2 and incubating at 30 °C to allow phase separation to occur, condensates are pelleted (P) by centrifugation and separated from soluble (S) proteins before gel analysis. D) 10 μM UBQLN2 (1% labeled with Alexa Fluor 647) was incubated ± 1 μM ubiquitinated substrate for 1 hr; Soluble (S) and pelleted (P) proteins were separated by centrifugation. E) UBQLN2 sedimentation as a function of substrate concentration. F) Quantification of replicate data (n=4-6) from E.

Mass spectrometry showed that the substrates were principally ubiquitinated on lysines within the degron region, although some substrates were also ubiquitinated within GFP, particularly in the mixed linkage substrate (**Supplemental Figure S1**). All substrates were highly ubiquitinated to a similar degree (estimated ≥15 Ub/substrate), with the bulk of the ubiquitinated substrate running at the top of the gel (**Figure 1B**).

### Ubiquitinated substrates condense with UBQLN2 in a linkage-specific manner

We first assessed the effect of ubiquitinated substrates on UBQLN2 phase separation via fluorescence microscopy (**Figure 1A**). Under conditions where full-length UBQLN2 exhibited little phase separation (10 μM UBQLN2, low salt, **Figure 1A**), K63-linked and mixed substrates (500 nM), but not K48-linked substrates, substantially promoted UBQLN2 phase separation (**Figure 1A,B**). Both UBQLN2 and polyubiquitinated substrates were localized into the same droplets, as previously observed for UBQLN2 and polyubiquitin chains (23).

We next quantified the effect of ubiquitinated substrates on UBQLN2 phase separation using sedimentation assays, in which components of the dense phase were pelleted upon centrifugation (**Figure 1C**). These sedimentation experiments additionally address the caveat that quantification of droplet imaging can be impacted by kinetics of droplet formation. Consistent with the microscopy data (**Figure 1A,B**), both K63-linked and mixed substrates (1 μM) induced robust phase separation of UBQLN2 (10 μM) while K48-linked substrates induced limited phase separation of UBQLN2 (**Figure 1D**). Similar results were obtained with 1 μM UBQLN2 (**Supplemental Figure S2**), in line with physiological concentrations (16). Phase separation was reciprocal (**Figure 1D**), such that ubiquitinated substrates co-sedimented with UBQLN2. There was a marked preference for longer chains in the condensates, as the highest molecular weight substrates were overrepresented in the pelleted fractions and underrepresented in the soluble fractions.

UBQLN2 sedimentation also depended on substrate concentration (**Figure 1E,F**). For K63-linked substrates, there was an increase in sedimentation with increasing substrate concentration. For K48-linked and, to a lesser extent, mixed substrates, biphasic behavior was observed, with an initial increase in sedimentation at lower substrate concentrations followed by a decrease at higher concentrations. Similar results were observed using enhanced GFP (eGFP) instead of sGFP (**Supplemental Figure S3**). These results agree with previous results using unanchored polyubiquitin chains, where K63-linked chains promoted UBQLN2 phase separation over a much broader concentration range than K48-linked chains (23). The biphasic behavior is consistent with re-entrant phase separation behavior stemming from heterotypic interactions between UBQLN2 and polyUb chains (23).

### Proteasomes localize into ubiquitinated substrate/UBQLN2 condensates

We next asked whether proteasome would follow the substrate into phase-separated condensates using sedimentation assays (**Figure 2, Supplemental Figure S4**). Incubating TagRFP-T-labeled proteasome (Methods) with Alexa647-labeled UBQLN2 and ubiquitinated substrates in the presence of proteasome peptidase inhibitors revealed that the proteasome significantly sedimented in the presence of UBQLN2 and both K63-linked and mixed-linkage substrates. There was also a trend towards increased proteasome sedimentation in the presence of the K48-linked substrate, although this was not statistically significant (p = 0.12), perhaps because under these conditions only some UBQLN2 phase separated (**Figure 1**). Interestingly, the proteasome’s deubiquitinase (DUB) activity, which was especially noticeable for the K63-linked substrate (31), was inhibited in the presence of UBQLN2 (compare substrate + UBQLN + proteasome to substrate + proteasome in **Figure 2A**). These data suggest that, even though both the substrate and proteasome are concentrated in the condensate pellet fraction, condensation may protect the substrate from proteasomal activity instead of enhancing it. Microscopy experiments confirmed co-localization of UBQLN2, proteasome and substrates (**Figure 2D**). Small proteasome puncta were observed in condensates regardless of linkage type (**Supplemental Figure S5**).

**Figure 2.**
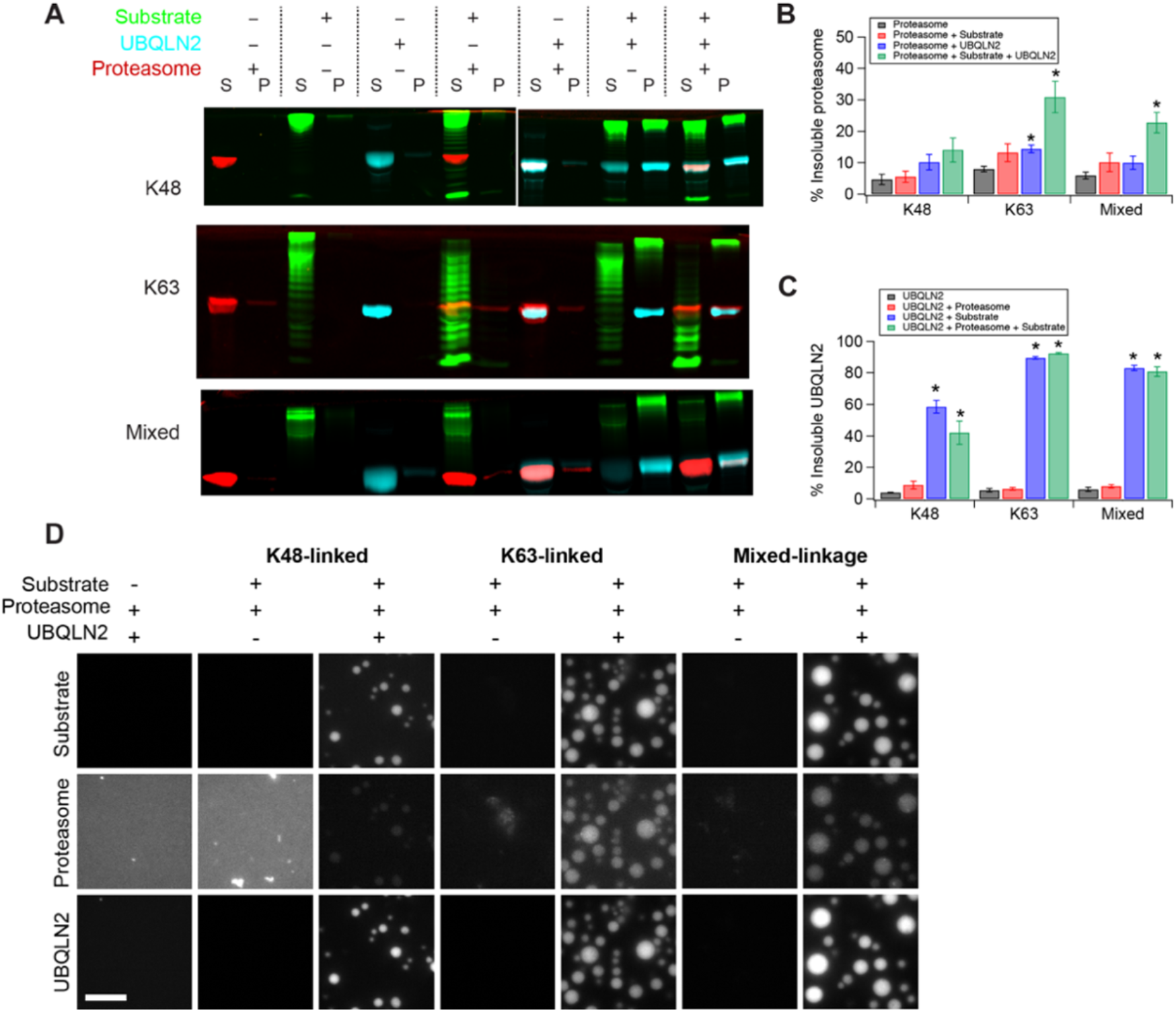
Proteasome is recruited to phase separated UBQLN2-substrate condensates. **A**) 100 nM TagRFP-T-Rpn6 containing proteasome and 500 nM K48-linked or K63-linked or 250 nM mixed linkage substrate were incubated with 10 μM UBQLN2 (1% Alexa 647 labeled) and 100 μM proteasome inhibitor cocktail (epoxomicin, MG132, bortezomib) for 1 hr as indicated. Individual channels are shown in **Supplemental Figure S4**. Soluble (S) and pelleted (P) proteins were separated by centrifugation and imaged using fluorescence. **B, C**) Quantification of replicate data from **A**. ^*^ indicates p*≤*0.05 relative to proteasome alone (**B**) or UBQLN2 alone (**C**) (two-tailed Welch’s t-test). Error bars are SEM from 3-4 measurements. **D**) Fluorescence microscopy of 10 μM UBQLN2, 100 nM proteasome, 500 nM ubiquitinated sGFP substrates, proteasome inhibitor cocktail, 1x deg buffer, and 1 hr at 30 ºC. Scale bar 10 μm.

### Substrates in UBQLN2 condensates can be protected from degradation

Does proteasomal recruitment to UBQLN2-substrate condensates have functional consequences? To test whether proteasome activity is enhanced (as has been shown for Rad23 in mammalian cells (13, 14)) or repressed, we examined the degradation of substrates in the presence of UBQLN2 condensates. We pre-incubated substrate and UBQLN2 to allow phase separation to occur, and then added proteasome to initiate the reaction. Since sGFP is quite stable, and, depending on ubiquitination state, is not always fully degraded by the proteasome (32, 33), we used a Cy5-labeled model substrate containing the same degron followed by two domains – the unstructured activation domain from the p160 transcriptional co-activator for thyroid hormone and retinoid receptors (ACTR) (34) followed by folded *E. coli* dihydrofolate reductase (DHFR) (35). Although DHFR is stably folded, the proteasome can easily unfold and degrade the substrate (with some small differences in the extent of degradation depending on chain type; red bars in **Figure 3B**) in the absence of UBQLN2 (**Figure 3**), as indicated by the Cy5-labeled peptides that appear at the bottom of the gel (35). UBQLN2 protected both K63-linked and mixed substrates, but not K48-linked substrates, against proteasomal degradation (**Figure 3A,B**, blue bars). One potential explanation would be that the substrates had different mobility within substrate/UBQLN2 condensates, leading to differential degradation rates. However, fluorescence recovery after photobleaching (FRAP) experiments demonstrated that K48- and K63-linked substrates exhibited similar mobility within condensates (**Supplementary Figure S7**). Another possible explanation would be that UBQLN2 binds ubiquitinated substrates and shields them from the proteasome. To decouple binding to ubiquitinated substrates from phase separation, we used a UBQLN2 deletion construct, the ΔSTI1-II mutant, which exhibits greatly reduced phase separation propensity (16, 18), but unaltered ability to bind to ubiquitin (**Supplemental Figure S8, Supplemental Table 1**). The presence of the ΔSTI1-II mutant failed to protect substrates from degradation, indicating that ubiquitin binding is not sufficient for protection, and that phase separation is required (**Figure 3A, B**, yellow bars). Crucially, under the same conditions as degradation occurs, the DHFR-containing substrate, like the sGFP substrate, reciprocally phase separates with UBQLN2, but not with the ΔSTI1-II mutant (**Figure 3C**).

**Figure 3.**
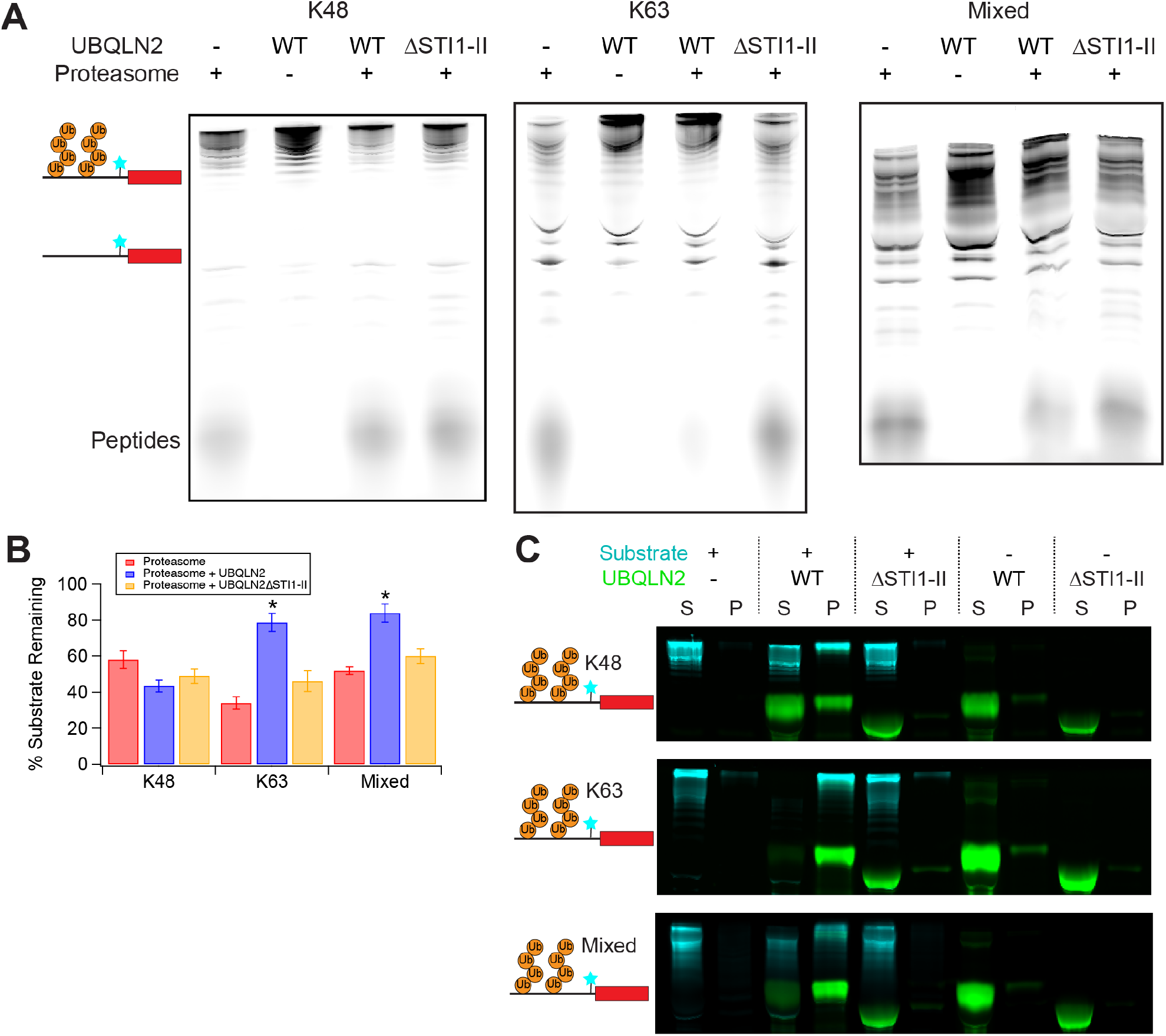
UBQLN2-induced phase separation inhibits proteasomal degradation in a polyUb-linkage dependent manner. **A)** Degradation of 250 nM ubiquitinated R-Neh2Dual-ACTR-DHFR substrate (Cy5-labeled N-terminal to DHFR) by 100 nM 26S proteasome with or without 10 μM UBQLN2 (1% DyLight-488 labeled) at 30°C for 30’. ΔSTI1-II is a UBQLN2 variant that does not phase separate on its own. **B)** Quantification of replicate data (% substrate remaining, quantifying total amount of full-length and ubiquitinated substrate) from **A**. Error bars are SEM from 3-4 measurements. * indicates p*≤*0.01 for proteasome + UBQLN2 relative to both proteasome alone and proteasome + UBQLN2ΔSTI1-II (two-tailed Welch’s t-test). Quantified regions of example gels are shown in **Supplemental Figure S6. C)** Sedimentation assays under the same condition as assays in **A** (but without proteasome) confirm reciprocal co-sedimentation of ubiquitinated substrates with WT but not ΔSTI1-II UBQLN2.

### Substrates in UBQLN2 condensates are protected from deubiquitination

Despite proteasomal recruitment to condensates, K63-linked and mixed substrates, but not K48-linked substrates, were protected from degradation and deubiquitination by the proteasome. To determine whether such protection was a general feature of UBQLN2-mediated condensates, we analyzed the ability of UBQLN2 to protect polyubiquitinated substrates from deubiquitination by DUBs. Ubiquitinated substrates and UBQLN2 were pre-incubated to allow condensate formation, and then different DUBs were added. We tested three DUBs: oTUB1, which is specific for K48-linked chains; AMSH, which is specific for K63-linked chains; and vOTU, which has no linkage specificity (36). Under conditions similar to those in which K63-linked and mixed substrates were protected from proteasomal degradation by UBQLN2, all three substrates (K48, K63 and mixed-linkage) were protected from DUB activity in the presence of WT UBQLN2 but not UBQLN2 ΔSTI1-II (**Figure 4A,B**). We therefore conclude that UBQLN2-mediated phase separation protects substrates from DUBs. Strikingly, K48-linked substrates show some protection from DUBs under the same conditions in which there is no protection from proteasomal degradation, indicating that there can be differential regulation of different enzymatic activities within condensates.

**Figure 4.**
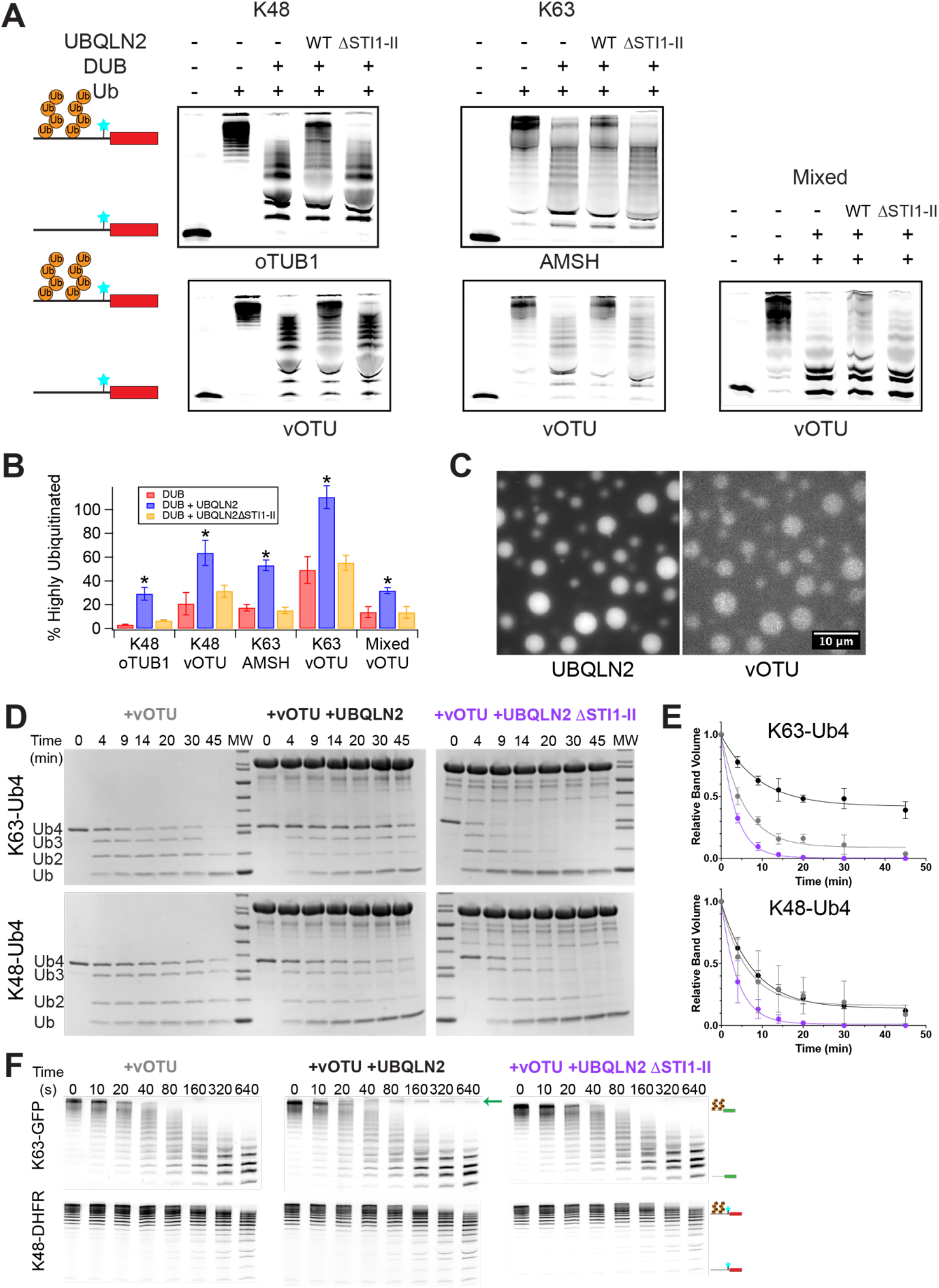
Phase separation protects polyubiquitin and polyubiquitinated substrates from DUB activity. **A)** Deubiquitination of 250 nM ubiquitinated (Ub +) R-Neh2Dual-ACTR-DHFR substrate (Cy5-labeled N-terminal to DHFR) by oTUB1, AMSH, or vOTU with or without 10 μM UBQLN2. Nonubiquitinated substrate (Ub -) is shown as a size reference. **B)** Quantification of replicate data (% highly ubiquitinated substrate remaining) from **A**. Error bars are SEM from 3-4 measurements. ^*^ indicates p<0.05 for DUB + UBQLN2 relative to both DUB alone and DUB + UBQLN2ΔSTI1-II (two-tailed welch’s t-test). Quantified regions of example gels are shown in **Supplemental Figure S9. C)** The linkage-independent DUB vOTU (100 nM AF488-labeled vOTU) colocalizes with 50 μM UBQLN2 in the presence of unlabeled 50 μM K63-Ub_4_. Scale bar = 10 μm. **D)** 25 μM Ub (6.25 μM Ub4) of K63-Ub_4_ or K48-Ub_4_ was incubated with 50 nM vOTU in the presence of 60 μM UBQLN2 or UBQLN2 ΔSTI1-II at 37 °C and time points were run on an SDS-PAGE gel. **E)** Quantification of DUB assays from **D**. The Ub_4_ band volumes were quantified as single-exponential fits (lines). Error bars are SD from n=3 DUB assay experiments. **F**) 100 nM each K48-linked R-Neh2Dual-ACTR-DHFR and K63-linked R-Neh2Dual-sGFP were incubated with 20 nM vOTU in the presence of 1 μM UBQLN2 or UBQLN2 ΔSTI1-II at 30 °C and time points were run on an SDS-PAGE gel and detected via fluorescence. Arrow indicates highly ubiquitinated K63-linked substrate that persists in the presence of UBQLN2.

Since different chain linkages have different abilities to induce phase separation, it might be possible to regulate DUB activity via differential phase separation. To explore this possibility, we used unanchored tetraubiquitin (Ub_4_), as K63-linked Ub_4_strongly induces UBQLN2 condensation while K48-linked Ub_4_represses UBQLN2 condensation (23). The linkage-independent DUB vOTU was recruited to condensates formed by K63-linked Ub_4_ and UBQLN2 (**Figure 4C**). The presence of WT UBQLN2 protected K63-linked Ub_4_ chains from disassembly by vOTU, but had no effect on K48-linked Ub_4_ (**Figure 4D,E**). Neither substrate was protected in the presence of the non-phase-separating ΔSTI1-II mutant, revealing that phase separation can act as a switch which converts a promiscuous DUB into a linkage-specific one. To further explore this possibility, we treated a mixture of K63-linked DHFR-containing substrate and K48-linked GFP-containing substrate with vOTU in the presence or absence of a physiological concentration (1 μM) of UBQLN2. Strikingly, while there was no substantial effect of UBQLN2 on deubiquitination of the K48-linked substrate, some of the highly ubiquitinated K63-linked substrate was protected from deubiquitination in the presence of UBQLN2 but not the ΔSTI1-II mutant (**Figure 4F**). Thus, in a complex cellular environment UBQLN2-dependent phase separation might protect a subset of ubiquitinated proteins.

## Discussion

Our results indicate that ubiquitinated substrates can induce phase separation of UBQLN2, and are themselves recruited to the UBQLN2 condensates (**Figure 5**). While homotypic interactions (e.g., oligomerization-driving STI1 interactions) can drive UBQLN2 to phase separate on its own, ubiquitinated substrates enable heterotypic interactions that further enhance UBQLN2 phase separation at lower, physiologically-relevant concentrations (**Supplemental Figure S2**). Consistent with previous results with unanchored polyUb chains (23), extended K63-linked substrates are the most potent drivers of phase separation, while compact K48-linked substrates induce some phase separation at lower substrate concentrations but then disfavor phase separation as the substrate concentration is increased. Branched chains containing multiple K48 and K63 linkages have intermediate properties. As there is little linkage-dependence to UBQLN2’s affinity for polyUb chains (23), these differences in phase separation behavior are likely driven by the different architectures of polyubiquitinated substrate-UBQLN2 complexes rather than by differences in affinity (24).

**Figure 5.**
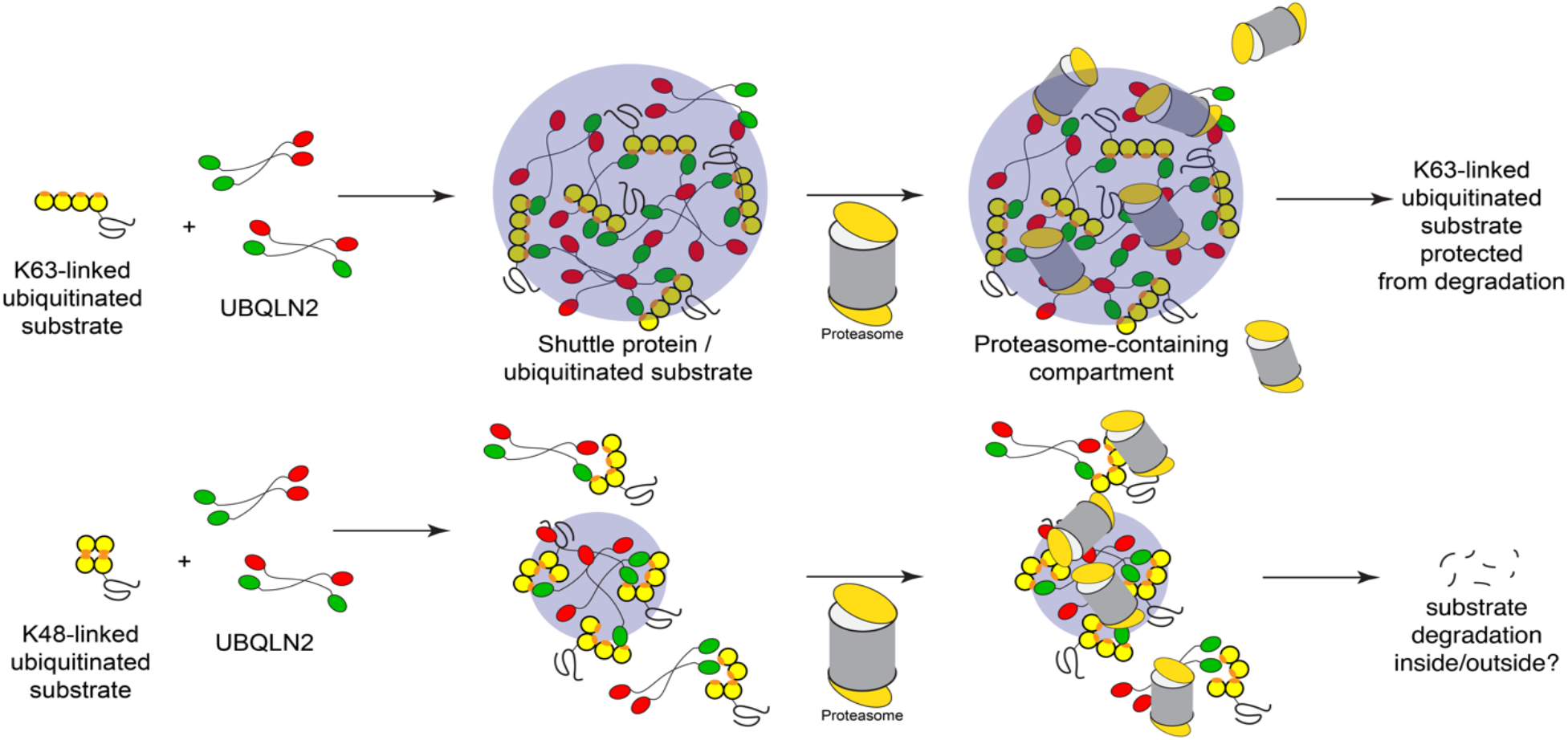
Ubiquitinated substrates condense with UBQLN2 to form membraneless compartments capable of recruiting protein quality control machinery. Our data show that ubiquitinated substrates can induce condensate formation with UBQLN2 as well as also be recruited into UBQLN2-substrate condensates, with K63-linked substrates showing greater propensity for this behavior over K48-linked substrates at similar substrate concentrations. Proteasomes can be recruited into these condensates although a proportion remains soluble and excluded from the condensate population. K63-linked ubiquitinated substrates are protected from degradation while K48-linked ubiquitinated substrates can still be degraded.

We observe a strong preference for longer chains in condensates, particularly for K63 and mixed-linkage chains. Conceptually, longer extended ubiquitin chains would be expected to serve as better scaffolds to enhance phase separation of shuttle proteins like UBQLN2, providing increased valency and multiple interaction surfaces (37), both via heterotypic Ub:UBA interactions and homotypic UBQLN2 STI1 interactions among other weaker interactions. We note that it is difficult to determine the physiological concentrations of highly ubiquitinated substrates. The substrate itself may be able to influence the phase separation process. However, the polyUb chains seem to be the major driver, as sGFP, eGFP and DHFR-containing substrates all showed qualitatively similar patterns of polyUb linkage-dependent phase separation. Similarly, although we cannot rule out a role for the location of the Ub modifications on the substrate in the phase separation process, there was no obvious correlation between predominant ubiquitination sites in the substrate and phase separation behavior. Moreover, although GFP can be directly ubiquitinated on surface lysines, the DHFR-containing substrate only contains lysines in the R-Neh2Dual degron, suggesting the site of ubiquitination is not controlling phase separation.

We find that UBQLN2-ubiquitinated substrate condensates, but not UBQLN2 or substrates on their own, recruit the proteasome. Interestingly, there appears to be some degree of internal structure to the condensates, with proteasome forming puncta within the larger UBQLN2-substrate droplets (**Supplemental Figure S5**). Puncta formation seems to be a pattern regardless of linkage type (K48 or K63), although we do note the appearance of some proteasome clustering even in the absence of UBQLN2 (**Figure 2D**). These results suggest the possibility for similar multicomponent condensates, potentially with internal organization, to form within the cell, although the condensates observed here *in vitro* are micron-sized, which are significantly larger than those observed in cells. Cellular condensates could also include other protein quality control components. For example, under various conditions UBQLN has been shown to interact with p62 and, separately, with various chaperones and other protein quality control components in nuclear condensates (38-40). In yeast, the UBQLN2 homolog Dsk2 interacts with both proteasomes and another shuttle factor Rad23 in nuclear condensates upon prolonged cell growth in a ubiquitin-dependent process (17).

Our data suggest that the proteasome is hindered from degrading substrates in the UBQLN2-substrate-proteasome condensates we form *in vitro*, at least for the K63-linked and mixed substates (**Figure 5**). There is a trend towards improved degradation of K48-linked substrates (p=0.052), but since these substrates incompletely induce phase separation, it is unclear whether this potential increase in proteasome activity is due to phase separation. Our results suggest an additional reason that, although K63- and K48-linked chains both lead to degradation *in vitro*, K63-linked chains are generally not used as proteasomal targeting signals *in vivo* (31, 41). In cells, additional components could potentially further modulate the fates of ubiquitinated substrates within condensates. For example, recruitment of ubiquitination machinery to condensates may also promote ubiquitination of substrates (42). Ubiquitination can be regulated within UBQLN2 condensates as UBQLN2 interacts with E3 ligases such as E6AP (43).

We suspect that the network architectures of UBQLN2-substrate phase separated condensates, governed by the interactions between UBQLN2 and ubiquitinated substrates with different polyUb chain linkages, can determine whether a substrate is protected or not from degradation or deubiquitination. Inclusion/exclusion of components may be regulated by the conformational entropy of disordered segments in proteins (44). Similarly, the differences in conformational ensembles of polyUb chains may play a role in regulating internal network architecture of condensates (24) and consequently, their activity. Intriguingly, our data show that, under the same conditions, all polyUb chain linkages are protected from DUB activity but only some linkages are protected from proteasomal degradation. Since both DUBs and the proteasome can be recruited to condensates, we presume that the effects of condensates on substrate protection are likely to be dependent on the substrate and cellular context. Although the exact mechanism of this differential protection remains unclear (*i*.*e*. whether degradation is occurring inside or outside of condensates), it points to the potential ability of phase separation to serve as a regulatory mechanism to control the fate of ubiquitinated substrates. This work adds to a growing body of literature that shows that phase separation has functional consequences, with condensates regulating the activity of multiple processes, including ubiquitination (42), immune system activation (45-47), and signaling in development (47, 48).

## Methods

### Constructs & Strains

Constructs expressing UBQLN2 and UBQLN2ΔSTI1-II (deletion of residues 379-462) were described previously (18). Constructs expressing His-SUMO-R-Neh2Dual-sGFP, His-SUMO-R-Neh2Dual-eGFP were cloned into previously existing His-SUMO-R-Neh2Dual constructs (27) using Gibson assembly and verified using Sanger sequencing. A construct expressing His-SUMO-R-Neh2Dual-ACTR-DHFR (which is lysine-free except for the degron region and contains a single cysteine between ACTR and DHFR) was described previously (35). An *S. cerevisiae* strain with TagRFP-T-Rpn6 (yDAK56) and a 3X-Flag-tag on Rpn11 was created using CRISPR as described previously (49).

### Protein Purification & Labeling

UBQLN2 proteins were expressed, purified and, as indicated, labeled with Alexa Fluor 647, Alexa Fluor 555, or DyLight-488 as described previously (16). Substrates were purified by Ni-NTA chromatography and SUMO protease cleavage as described previously, and R-Neh2Dual-ACTR-DHFR was labeled with sulfo-cyanine5 maleimide (Lumiprobe) on a single unique cysteine immediately N-terminal to DHFR, and repurified by gel filtration as described previously (27, 35). 26S Proteasome was purified from *S. cerevisiae* via a 3X-FLAG tag on Rpn11 as described previously (27). AMSH*, oTUB1* and vOTU were purified as described previously (33). vOTU was labeled with Alexa Fluor 555 and repurified by gel filtration. Ub_4_ was synthesized and purified as described previously (23). Substrate, UBQLN2 and DUB protein sequences are given in **Supplemental Table S2**.

### Ubiquitination

Substrates were ubiquitinated largely as described previously (27). To generate K63-linkages, substrates (5 μM) were incubated with mammalian E1 (166 nM), UbcH7 (E2, 2.9 μM), and Rsp5 (E3, 2.9 μM) ligases in 25 mM Tris-Cl pH 7.5, 50 mM NaCl, 4 mM MgCl_2_, 1.33 mg mL^-1^ ubiquitin, 4 mM ATP, and 1 μM DTT. To generate K48-linkages, substrates (5 μM) were incubated with mammalian E1 (100 nM), Ubc2 (E2, 12 μM), and Ubr1 (E3, 400 nM) ligases in 50 mM HEPES pH 7.5, 150 mM NaCl, 10 mM MgCl_2_, 1.33 mg mL^-1^ ubiquitin, and 5 mM ATP. To generate mixed linkages, substrates (4 uM) were incubated with mammalian E1 (E1, 130 nM), UbcH5 (E2, 4 μM), Cul3/Rbx1 (E3, 4 μM) and Keap1 C151S (4 μM) in 45 mM Tris-Cl pH 8.0, 100 mM NaCl, 10 mM MgCl_2_, 0.73 mg mL^-1^ ubiquitin, 2 mg mL^-1^ ovalbumin and 5 mM ATP. All reactions took place in the dark at 30°C for 1.5-2 hours. Polyubiquitinated substrates were purified in 1x degradation assay buffer (50 mM Tris-Cl pH 7.5, 5 mM MgCl_2_, 5% glycerol) by spin size exclusion using Sephadex G75. Concentrations were determined using the fluorescence intensity of the final sample.

### Mass spectrometry

On-column digests of ubiquitination reactions were performed using the Protifi S-Trap™ mini kit (K02-Mini-10) following the manufacturer’s protocol in preparation for peptide analysis by LC-MS. The proteases trypsin and Lys-C (Promega, V5073) were used to digest the LtEc proteins overnight at 37°C after reduction with 5 mM TCEP at 55°C and alkylation with 20 mM iodoacetamide at room temperature. Peptides were separated on an Agilent InfinityLab Poroshell 120 EC-C18 (50 x 2.1 mm, 2.7 μm particle, # 699775-902) column maintained at 40°C on a SCIEX Exion LC (Framingham, MA). The HPLC solvents used were (A) water with 0.1% formic acid and (B) acetonitrile with 0.1% formic acid at a flowrate of 0.4 mL/min. Gradient elution consisted of 100% A hold for 5 minutes, followed by two gradient steps (0-40% B from 5-60 min and 40-95% B from 60-75 min) with a final hold at 95% B from 70-90 minutes. Eluted peptides were analyzed by data dependent acquisition (DDA) on a SCIEX 5600+ TripleTOF mass spectrometer calibrated with the SCIEX APCI positive calibrant solution prior to each analysis. Positive ESI precursor ions for DDA were determined under high sensitivity mode TOF from 200–1250 Da with the following source parameters: DP = 100 V, CE = 10, GAS1 = GAS2 = 60 psi, CUR = 30 psi, ISFV = 5500 V, and a source temperature of 500°C. DDA fragmentation experiments selected the ten most abundant ions from survey scan criteria between 400-1250 m/z, with charge states between 2+ to 5+, using dynamic background selection and rolling collision energy (CE). Selected target ions were excluded for 7 sec (2 cycles). Spectra were analyzed for ubiquitin modification using ProteinPilot (SCIEX, v. 5.0.2), with a custom database consisting of the proteins known to be present in the ubiquitination reactions (E1, E2, E3 enzymes, substrate and ubiquitin). A custom modification to tyrosine (loss of OH4) was used to enable detection of the GFP chromophore derived from TYG; false positives containing this modification on other peptides were discarded. Distinct peptides at an FDR of 1% (or, for Keap1 ubiquitination, at a confidence level of 99%) were used to assess locations of ubiquitin modification, with manual assignments of ambiguous peptides based on theoretical confidence levels and the typical absence of a cleavage event after a ubiquitinated lysine. Data is deposited as MassIVE MSV000093780.

### Sedimentation assays

UBQLN2 was incubated with polyubiquitinated substrates in 1X degradation assay buffer containing 2 mM DTT, 1 mg mL^-1^ BSA, and 1% DMSO in Eppendorf Protein Lo-Bind tubes for 1 hour at 30°C. For titration experiments, substrates were serially diluted in 1x degradation assay buffer prior to the addition of UBQLN2. For experiments including TagRFP-T-Rpn6 proteasome, reaction mixture also contained 1 mM ATP, 10 mM creatine phosphate, 0.1 mg mL^-1^ creatine phosphokinase and a proteasome inhibitor cocktail consisting of 100 μM each epoxomicin, bortezomib, and MG132.

Following incubation, reactions were spun at 15871 x *g* for 5 minutes at room temperature to separate soluble and phase-separated proteins. Pellets were solubilized in 2X SDS-PAGE sample buffer (50). Proteins were run on 9.25% tris-tricine gels, scanned on a Typhoon FLA 9500 imager (Cytiva) with voltage adjusted to assure all samples were within the linear range of the detector, and relative amounts of UBQLN2, substrate, and proteasome present in soluble and pelleted fractions were determined using ImageQuant.

### Microscopy

Samples were prepared on ice to contain either 10 μM UBQLN2 (spiked with 20 nM UBQLN2 labelled with Alexa Fluor 647) and/or 500 nM of sGFP substrates in 1x degradation assay buffer. Samples with 100 nM TagRFP-T-Rpn6 containing proteasome were also prepared with 100 μM proteasome inhibitor cocktail. Samples were added to Eisco Labs Microscope Slides, with Single Concavity, and covered with MatTek coverslips that had been coated with 5% bovine serum albumin (BSA) to minimize changes due to surface interactions, and incubated coverslip-side down at 30°C for 1 hour. Samples of deubiquitinase vOTU contained 50 μM UBQLN2 (spiked with Alexa Fluor 647-labelled UBQLN2), 50 μM Ub of K63-Ub4 (spiked with Alexa Fluor 488-labeled K63-Ub4), 20 mM HEPES, 200 mM NaCl, 1 mM DTT, pH 7, and were incubated at 30 ºC and imaged at indicated time points. Phase separation was imaged on an ONI Nanoimager (Oxford Nanoimaging Ltd, Oxford, UK) equipped with a Hamamatsu sCMOS ORCA flash 4.0 V3 camera using an Olympus 100×/1.4 N.A. objective. Images were prepared using Fiji (51) and FigureJ plugin.

### Microscopy analysis

For puncta count of droplet area, image background subtraction was performed using a rolling ball radius of 50 pixels. Images were subsequently thresholded using default settings. Puncta were analyzed using a minimum size of 10 pixel^2 and a circularity range of 0.3 - 1.0 to eliminate edge artifacts. To address the important caveat that kinetics of droplet formation may impact droplet area, we also measured the insoluble fraction of UBQLN2 at different concentrations of substrate (see Figure 1F and Supplemental Figure S3D). The insoluble fraction measurements are indirect indicators of the saturation concentration (c_sat_) of UBQLN2 typically found in multi-component phase diagrams.

### Fluorescence Recovery After Photobleaching (FRAP)

Samples were prepared to contain 10 μM UBQLN2 and 125 nM of either K48-or K63-linked substrates in 1X degradation assay buffer. Samples were added to Eisco Labs Microscope Slides, with Single Concavity, and covered with MatTek coverslips that had been coated with 5% bovine serum albumin (BSA), and incubated coverslip-side down at 37 °C for 20-30 min. FRAP was carried out on a Zeiss LSM 980 with Airyscan 2 confocal microscope (Carl Zeiss AG, Oberkochen, Germany) using a Plan-Apochromat 63X/1.4 NA oil. Images were prepared using Fiji (51) and FigureJ plugin.

### Degradation assays

Cy5-labeled polyubiquitinated R-Neh2Dual-ACTR-DHFR (250 nM) was preincubated with 10 μM UBQLN2 in 1x degradation assay buffer containing 1 mM ATP, 10 mM creatine phosphate, 0.1 mg mL^-1^ creatine phosphokinase, 2 mM DTT, 1 mg mL^-1^ BSA, and 1% DMSO for 1 hour at 30°C. 100 nM 26S proteasome was then added, and the reaction was incubated for anther 30 minutes. Reactions were stopped by the addition of an equal volume of SDS-PAGE sample buffer (100 mM Tris pH 6.8, 24% glycerol, 8% SDS, 0.2 mg mL^-1^ coomassie blue R250, 0.2 M DTT). Samples were run on 9.25% tris-tricine gels, scanned on a Typhoon FLA 9500 imager with voltage adjusted to assure all samples were within the linear range of the detector, and relative amounts of substrate remaining in each reaction were determined using ImageQuant.

### Deubiquitinase assays

*Using polyubiquitinated substrates*: Cy5-labeled polyubiquitinated R-Neh2Dual-ACTR-DHFR (250 nM) was preincubated with 10 μM UBQLN2 in 1x degradation assay buffer containing 2 mM DTT, 1 mg mL^-1^ BSA, and 1% DMSO for 1 hour at 30°C. For K63-linked substrates, 1 μM AMSH or 40 nM vOTU was added and the reactions were incubated for 15 or 10 min, respectively. For K48-linked substrates, 1 μM OTUB1 or 200 nM vOTU was added and the reactions were incubated for 20 or 10 minutes, respectively. For mixed linkage substrates, 200 nM vOTU was added and the reaction was incubated for 30 minutes. Incubation times and DUB concentrations were chosen in control experiments to ensure partial deubiquitination in the absence of UBQLN2. DUB reactions were stopped by the addition of an equal volume of SDS-PAGE sample buffer. Samples were run on 9.25% tris-tricine gels, scanned on a Typhoon FLA 9500 imager with voltage adjusted to assure all samples were within the linear range of the detector, and relative amounts of highly ubiquitinated substrate remaining in each reaction were determined using ImageQuant.

### Using free polyUbiquitin chains

Samples were prepared on ice to contain 50 nM vOTU, 30 μM of K48-Ub4 or K63-Ub4, with and without 60 μM of full-length UBQLN2 or UBQLN2 ΔSTI1-II in 20 mM NaP, 500 mM NaCl, pH 6.8. Samples were aliquoted into separated microfuge tubes and incubated at 37 ºC. At indicated time points, the reactions were stopped by the addition of an equal volume of SDS-PAGE sample buffer. Samples were run on 4-20% gradient gels (BioRad), stained with Coomassie Blue, destained, imaged on Gel Doc EZ imager (BioRad) and bands intensities were quantified with Image Lab (BioRad).

### Nuclear Magnetic Resonance (NMR) Spectroscopy

Protein solutions of 50 μM ^15^N UBQLN2 or UBQLN2 ΔSTI1-II were prepared in 20 mM NaPhosphate buffer (pH 6.8) with 0.5 mM EDTA, 0.02 % NaN3, and 5 % D_2_O. NMR data (either standard ^1^H-^15^N SOFAST-HMQC or ^1^H-^15^N TROSY-HSQC experiments) were acquired at 25°C using a Bruker Avance III 800 MHz spectrometer equipped with TCI cryoprobe. NMR data were subsequently processed using NMRPipe (52) and analyzed using CCPNMR 2.5.2 (53). UBA peak assignments were transferred by visual inspection using previously-determined UBQLN2 450-624 assignments (16). To determine Ub binding affinity (*K*_d_), we prepared separate NMR samples with known amounts of unlabeled Ub (up to a stoichiometry of ∼2:1 Ub:UBQLN2). Binding was monitored as a function of different ligand:protein ratios. Assuming the fast-exchange limit, the chemical shift perturbation (CSP) for each backbone amide was calculated as 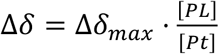, where Δδ_*max*_ is the CSP in the fully-bound state, and [PL] and [Pt] represent the ligand-bound and the total UBQLN2 protein concentrations, respectively. The CSP (Δδ) is quantified as 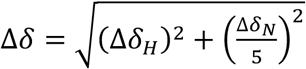 where Δδ_*H*_ and Δδ_*N*_ are the differences in ^1^H and ^15^N chemical shifts in ppm for amide resonances compared between the NMR spectrum at a specific titration point and the ligand-free NMR spectrum. Titration data for each amide resonance were fit to the single-site binding model with an in-house MATLAB program: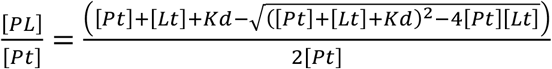.[Lt] is the total ligand (Ub) concentration and *K*_d_ is the binding affinity. Reported *K*_d_ values are averages of residue-specific *K*_d_ values with the error reflecting the standard deviation of these values.s

## Funding

This material is based upon work supported by the National Science Foundation under Grant No. 1935596 to D.A.K (sedimentation assays, substrates, proteasome, DUBs). This work was supported by NIH R01GM136946 (UBQLN2 protein purifications, Ub_4_, microscopy, NMR) to C.A.C.. FRAP experiments were performed on a Zeiss LSM980 with Airyscan2 supported by NIH S10 OD026946-01A1. Data collected on a Bruker 800 MHz NMR magnet (SU, SUNY-ESF) was supported by NIH shared instrumentation grant 1S10OD012254. Data collected on a SCIEX 5600+ was supported by a Major Research Instrumentation grant from the National Science Foundation (Grant No. CHE-2018399).

## Acknowledgements

The authors thank members of the Kraut and Castañeda labs, especially Dr. Edwin Ragwan, Dr. Billy Haws, Dr. Anitha Rajendran, Dr. Nirbhik Acharya, as well as Dr. Jeroen Roelofs and Dr. Jeremy Schmit for valuable discussions and Matt Schnell, Dr. Anthony Lagalante and Luke Lagalante for assistance with mass spectrometry experiments. We acknowledge additional microscopy support from the SU Bioimaging Center.

## Data Availability

Mass spectrometry data for ubiquitinated substrates are deposited in MassIVE: ftp://massive.ucsd.edu/v06/MSV000093780/. All other data are available online via DOI 10.25833/tcra-t419.

## Notes

ftp://massive.ucsd.edu/v06/MSV000093780/

### Competing Interest Statement

The authors have declared no competing interest.

### Summary of Updates

Textual revisions to improve clarity, additional FRAP experiments in supplementary materials.

