## Supplemental Information for "Phase separation of polyubiquitinated proteins in UBQLN2 condensates controls substrate fate"

Isabella M. Valentino[1], Jeniffer G. Llivicota-Guaman[1], Thuy P. Dao[2], Erin O. Mulvey[1], Andrew M. Lehman[1], Sarasi K. K. Galagedera[2], Erica L. Mallon[1], Carlos A. Castañeda[2]*, Daniel A. Kraut[1]*
*co-corresponding authors

[1]Department of Chemistry, Villanova University, Villanova, PA 19085
[2]Department of Biology, Department of Chemistry, Bioinspired Institute, Interdisciplinary Neuroscience Program, Syracuse University, Syracuse, NY 13244

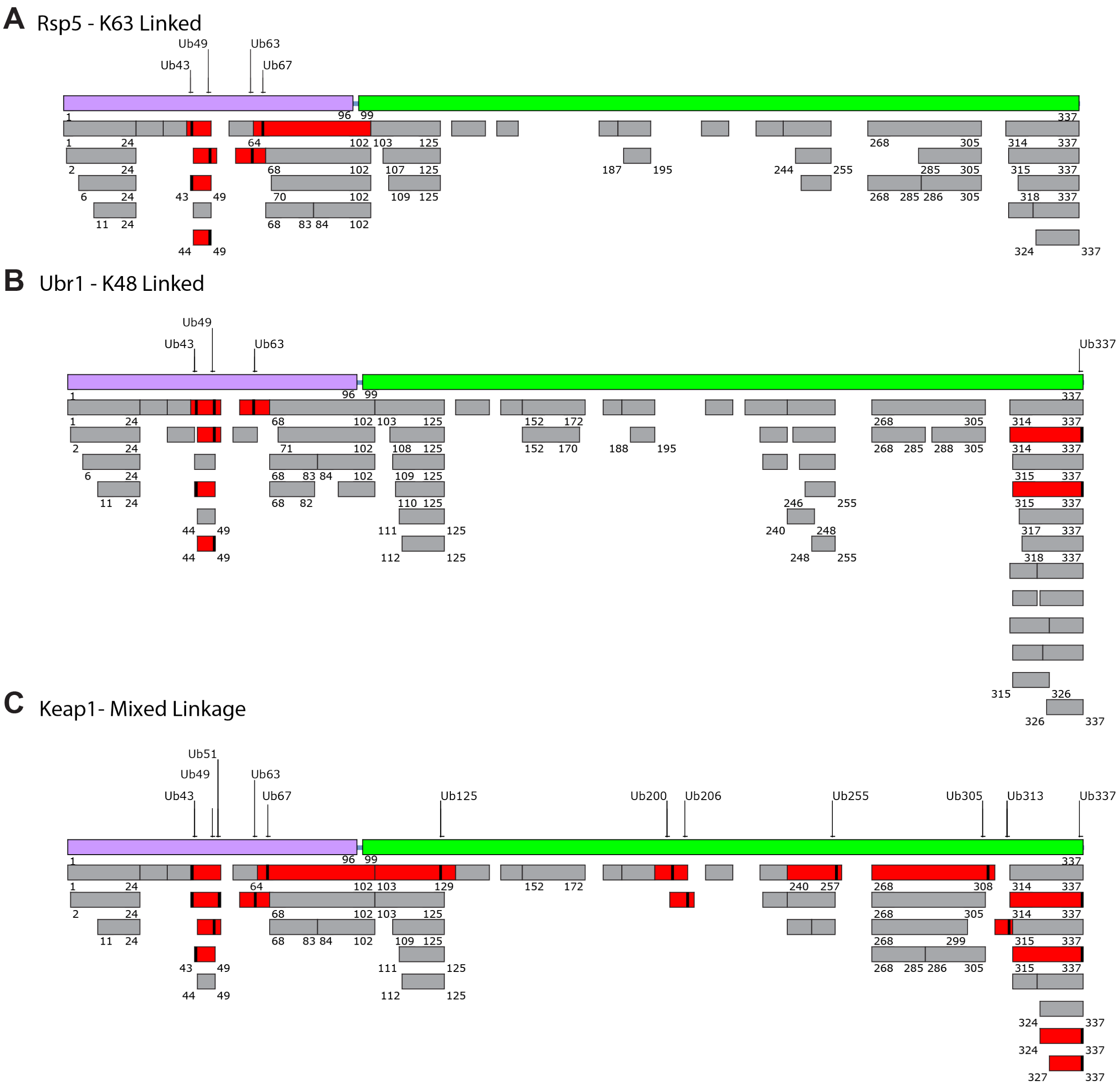


**Supplemental Figure S1**. Mass spectrometry analysis of **A)** Rsp5-ubiquitinated (K63-linked), **B)** Ubr1-ubiquitinated (K48-linked) and **C)** Cul3/Rbx1/Keap1-ubiquitinated (mixed linkage) substrates. Protein sequence map shows R-Neh2Dual degron in purple, sGFP in green, and observed sites of ubiquitin modification as indicated. Unique tryptic peptides are shown below the map. Those with evidence for ubiquitin modification (GG or LRGG adducts) are shown in red with a black line at the site of modification, while non-ubiquitinated peptides are shown in grey.

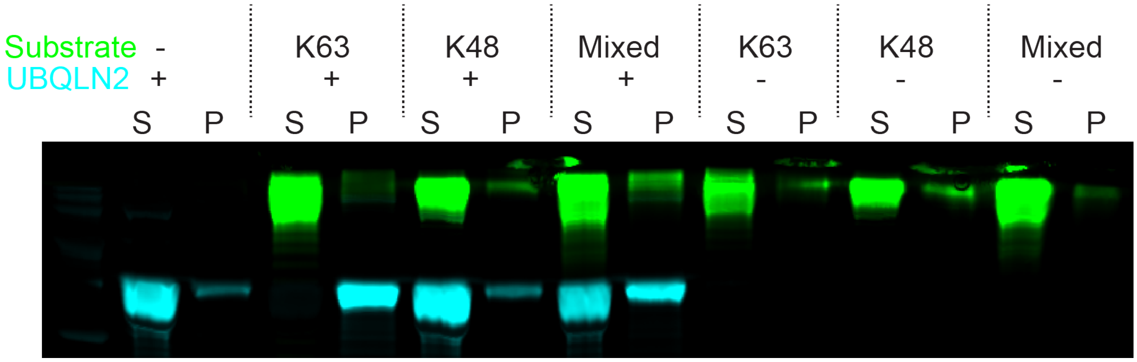


**Supplemental Figure S2**. K63 and mixed linkage chains sediment UBQLN2 at physiological UBQLN2 concentrations. 1 μM UBQLN2 (1% labeled with Alexa Fluor 647) was incubated ± 1 μM ubiquitinated Neh2Dual-sGFP substrate for 1 hr; soluble (S) and pelleted (P) proteins were separated by centrifugation.

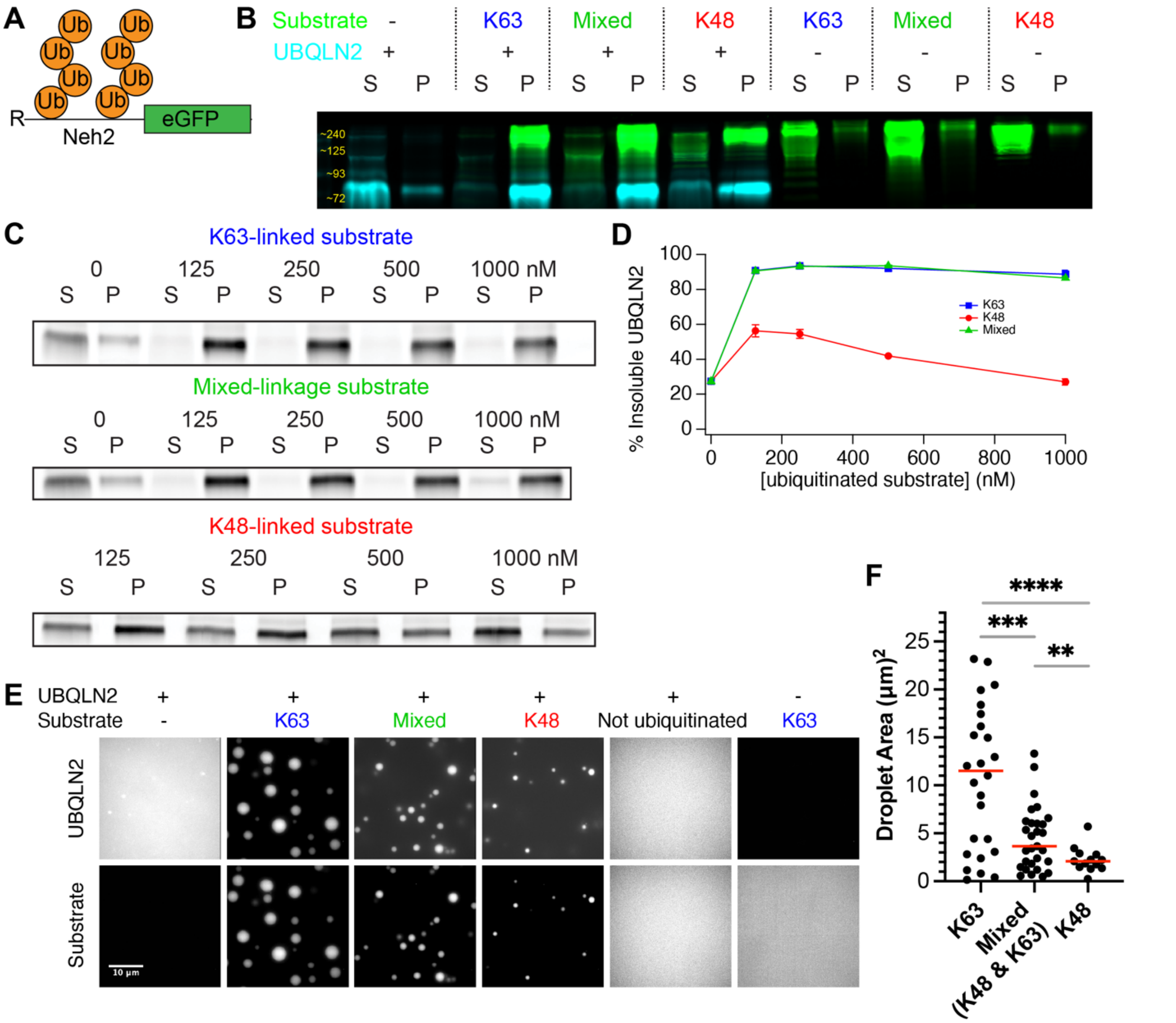


**Supplemental Figure S3.** Ubiquitinated eGFP substrate sediments UBQLN2 in a linkage-dependent manner. A) Assays use R-Neh2Dual-eGFP substrate ubiquitinated with K63-linked (Rsp5), K48-linked (Ubr1), or mixed linkage chains (Keap1/Cul3/Rbx1). B) 10 µM UBQLN2 (1% labeled with Alexa Fluor 647) was incubated ± 250 nM ubiquitinated substrate for 1 hr; soluble (S) and pelleted (P) proteins were separated by centrifugation. C) UBQLN2 sedimentation as a function of substrate concentration. D) Quantification of replicate data from C. Error bars are SEM from 3-4 measurements. E) Fluorescence microscopy of 10 µM UBQLN2 (1% labeled with Alexa Fluor 647) with or without 250 nM ubiquitinated (or not Ub'ed) substrate for 15 minutes at 30 ºC. F) Quantification of droplet size from images in E (K63: n=26, Mixed: 29, K48: 13 droplets) and statistics applied with Welch's two-tailed t-test (****, $p < 0.0001$; ***, $p < 0.001$; **, $p < 0.01$).

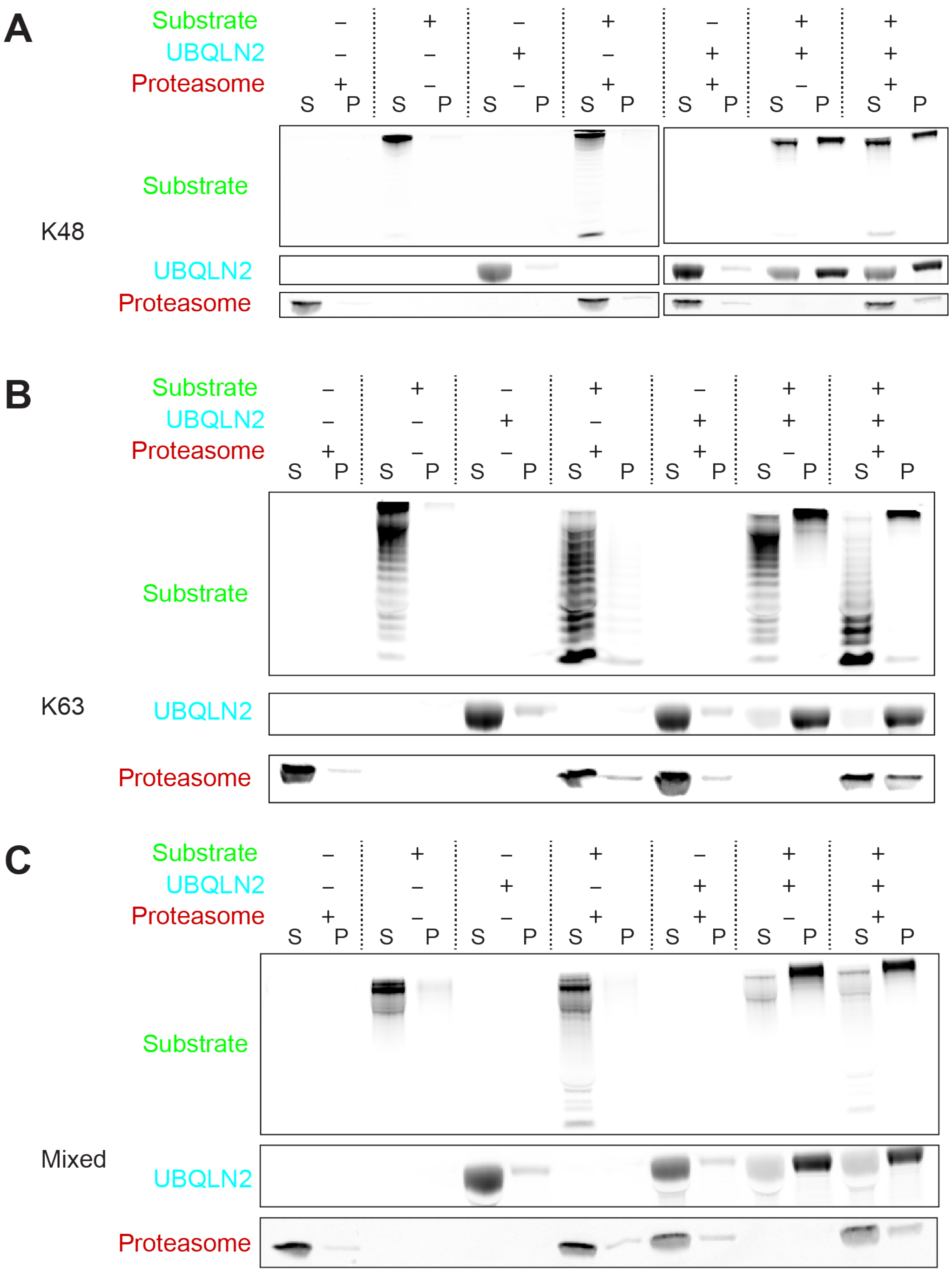


**Supplemental Figure S4**. Individual fluorescence channels from **Figure 2A**.

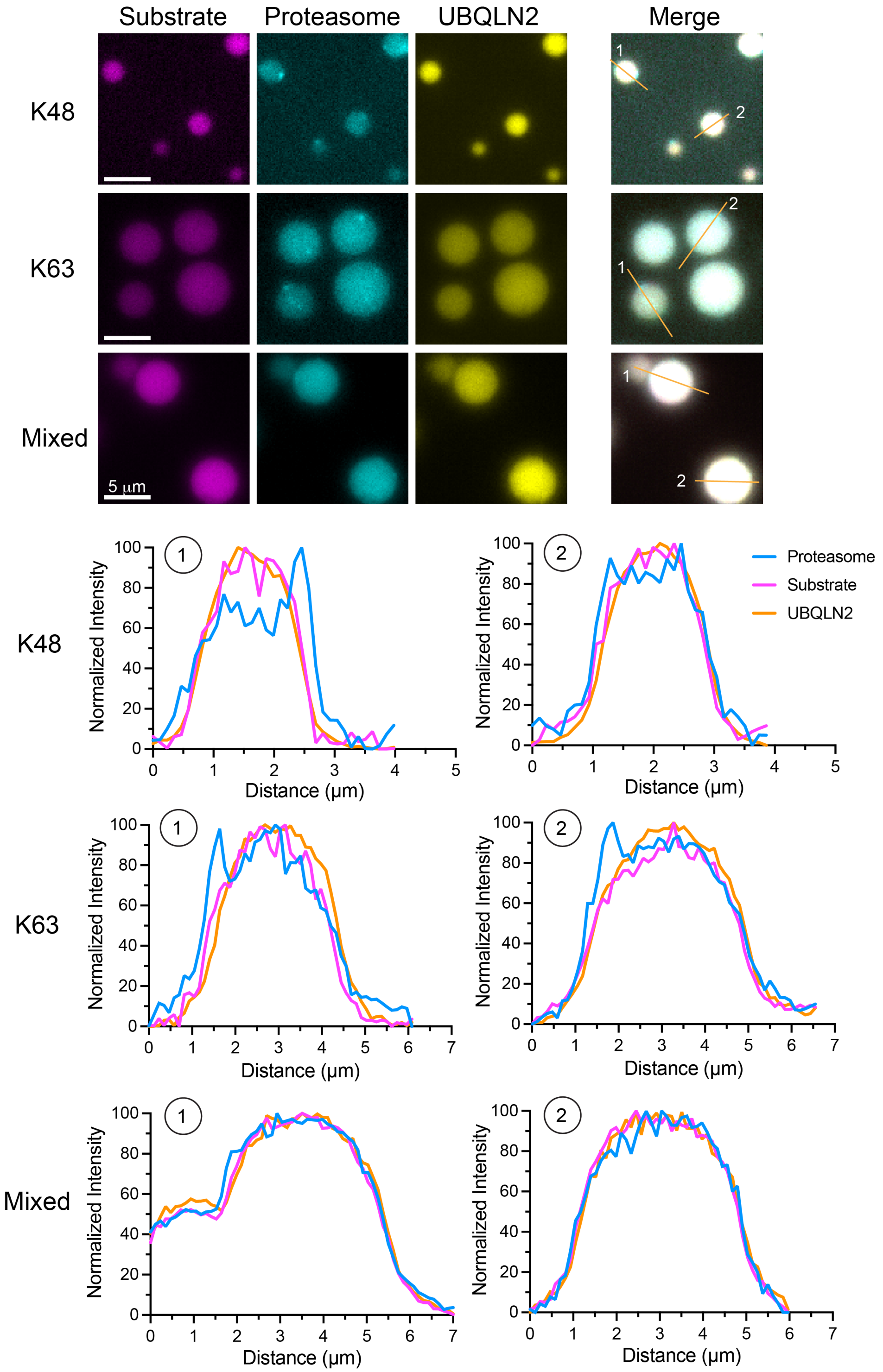


**Supplemental Figure S5.** Representative substrate-UBQLN2 condensates containing proteasome puncta. Contrast was adjusted on images to highlight puncta within condensates. Line traces are shown for two condensates in each set of images. Scale bar is 5 μm.

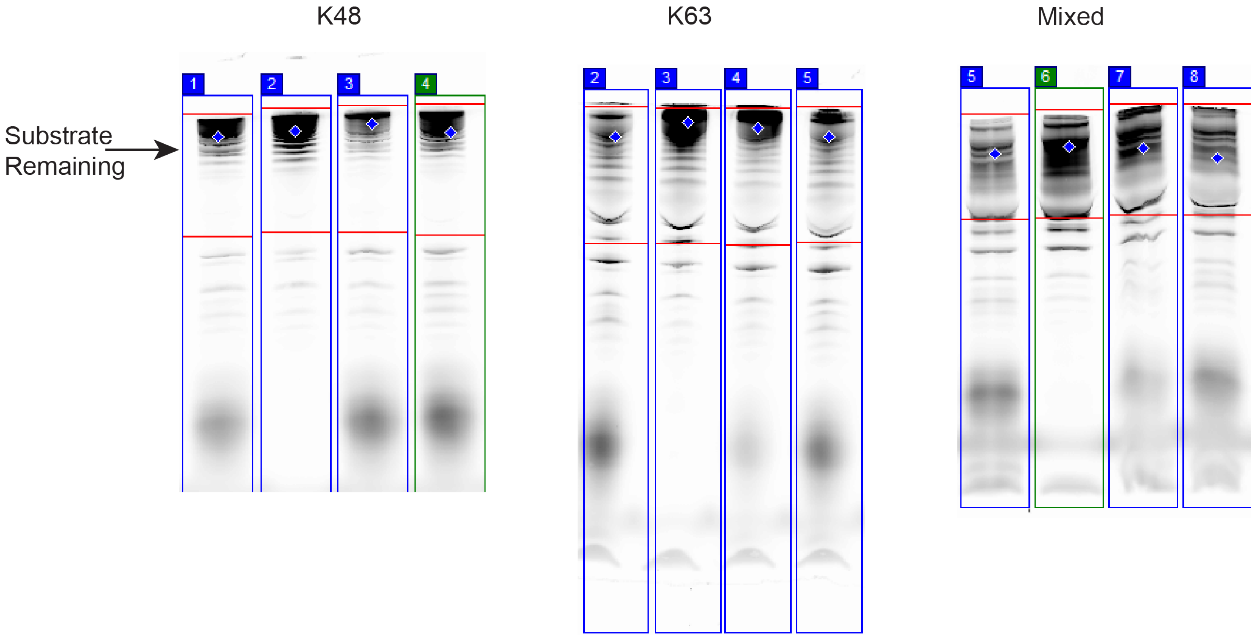


**Supplemental Figure S6**. **Quantification of sample gels from Figure 3A.** Red boxed area at the top shows the approximate areas that were quantified to determine the extent of degradation (see arrow); full-length non-ubiquitinated substrate was included in the quantification to ensure deubiquitination was not misinterpreted as degradation.

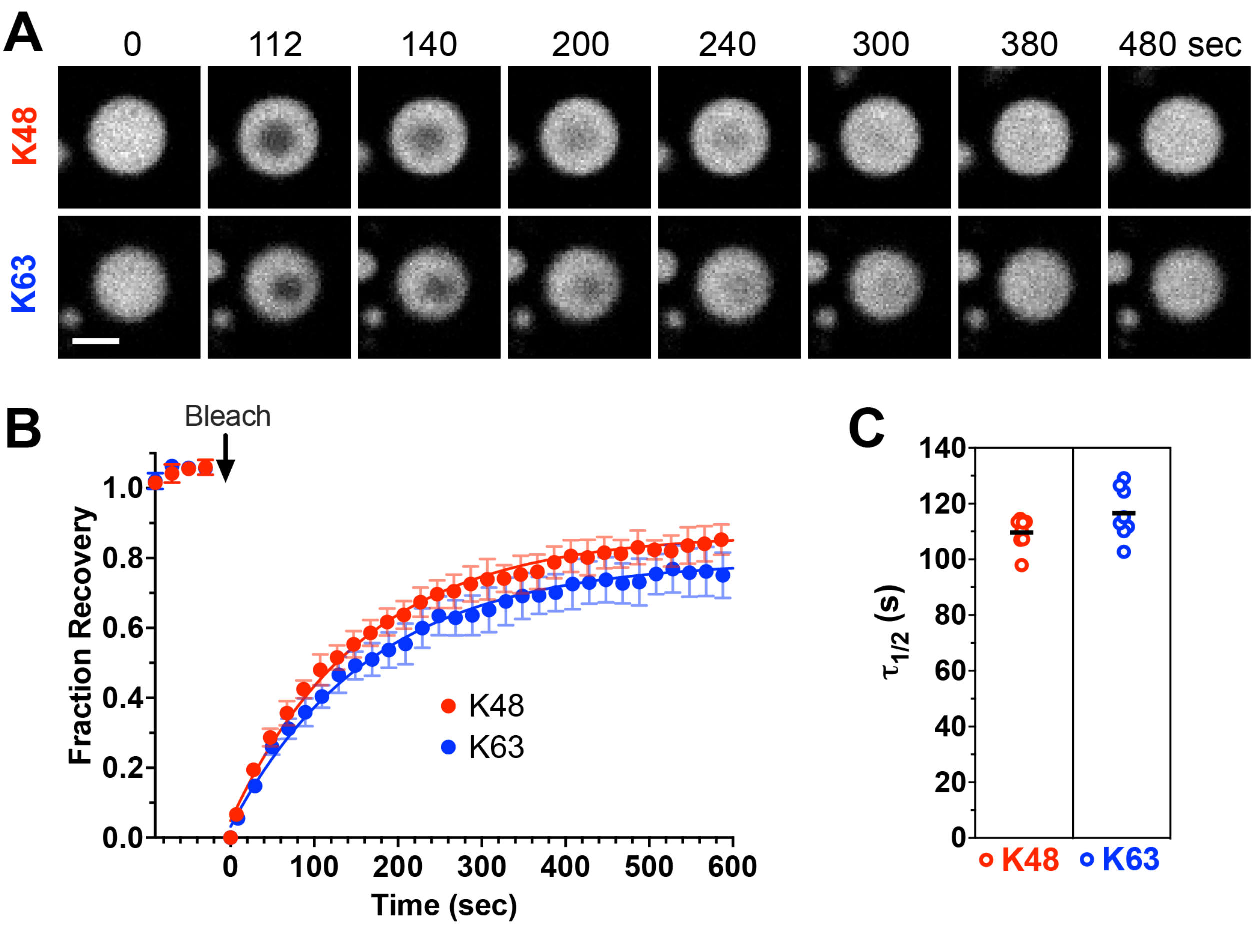


**Supplemental Figure S7. A)** Representative fluorescence images of partial droplet photobleaching experiments for K48- or K63-linked ubiquitinated R-Neh2Dual-sGFP substrates in UBQLN2/substrate droplets at 125 nM substrate and 10 µM UBQLN2 concentrations. Scale bar is 10 µm. **B)** Normalized fluorescence intensity over time for droplets. Dots indicate the average data points from 8 separate droplets. Lines indicate a single exponential fit to the data. Error bars represent the SD. **C)** Half-times of fluorescence recovery of K48 or K63 substrates in UBQLN2/substrate droplets. Black lines represent the mean.

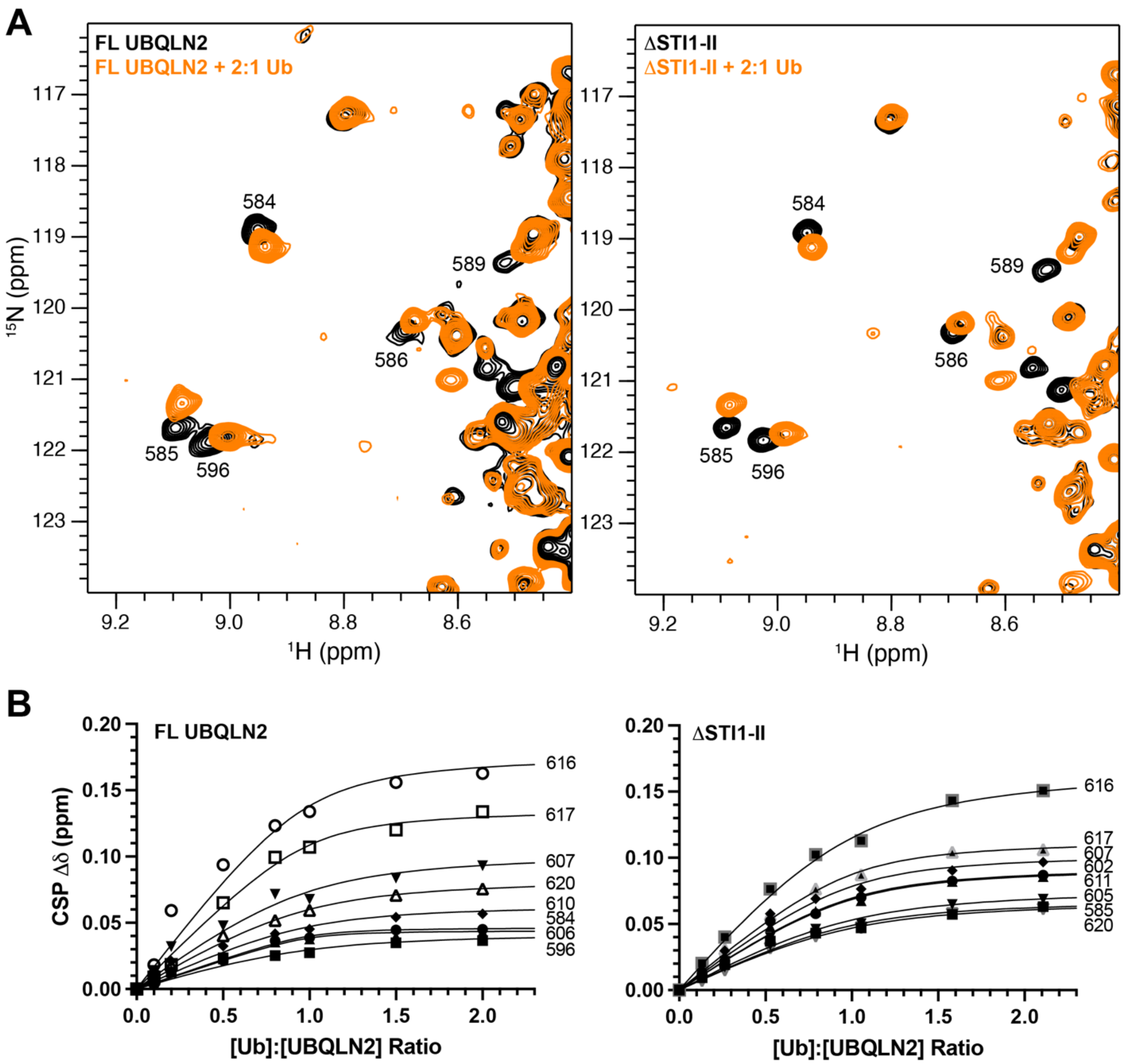


**Supplemental Figure S8. NMR analysis of full-length UBQLN2 and ΔSTI1-II in the presence of Ub. A)** $^{1}$H-$^{15}$N TROSY-HSQC NMR spectra of either 50 μM (left) full-length UBQLN2 or (right) ΔSTI1-II in the absence (black) and presence (orange) of Ub (~2:1 Ub:UBQLN2 stoichiometric ratio). **B)** Amide titration curves for select UBA resonances (labeled) with lines representing fits of single-site binding model to titration data (see Methods and Supplemental Table 1). Data for FL UBQLN2 are identical to those presented in {Dao et al., 2022, #267227}.

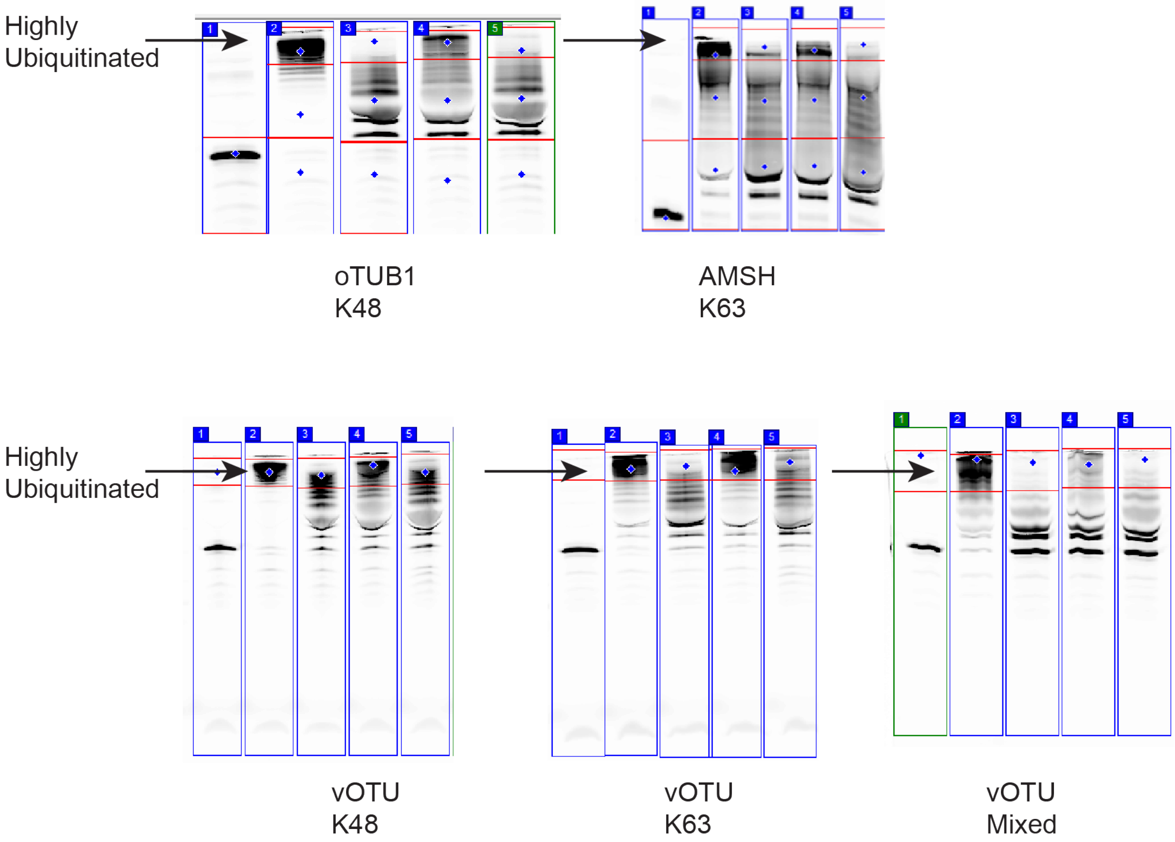


**Supplemental Figure S9**. **Quantification of sample gels from Figure 4A.** Red boxed area at the top (see arrow) shows the approximate areas that were quantified to determine the extent of highly ubiquitinated substrate remaining.

**Supplemental Table S1.** Residue-specific titration curves as determined from NMR spectroscopy.

| **FL** | | **ΔSTI1-II** | |
|---|---|---|---|
| Res # | $K_D$ (μM) | Res # | $K_D$ (μM) |
| 584 | 1.1 | 585 | 5.8 |
| 596 | 6.7 | 602 | 4.8 |
| 606 | 0.9 | 605 | 7.2 |
| 607 | 5.7 | 607 | 3.8 |
| 610 | 4.6 | 611 | 4.5 |
| 616 | 3.0 | 616 | 7.5 |
| 617 | 2.4 | 617 | 3.4 |
| 620 | 5.4 | 620 | 7.2 |
| Avg | 4 ± 2 | Avg | 6 ± 2 |

For FL UBQLN2, $K_D$ is 3.7 ± 2.2 μM; For ΔSTI1-II, $K_D$ is 5.5 ± 1.6 μM

**Supplemental Table S2**. Sequences of substrates and UBQLN2 constructs.

| Protein | Sequence |
|---|---|
| UBQLN2 | MAENGESSGPPRPSRGPAAAQGSAAAPAEPKIIKVTVKTPKEKEEF<br>AVPENSSVQQFKEAISKRFKSQTDQLVLIFAGKILKDQDTLIQHGIHD<br>GLTVHLVIKSQNRPQGQSTQPSNAAGTNTTSASTPRSNSTPISTNS<br>NPFGLGSLGGLAGLSSLGLSSTNFSELQSQMQQQLMASPEMMIQI<br>MENPFVQSMLSNPDLMRQLIMANPQMQQLIQRNPEISHLLNNPDIM<br>RQTLEIARNPAMMQEMMRNQDLALSNLESIPGGYNALRRMYTDIQE<br>PMLNAAQEQFGGNPFASVGSSSSSGEGTQPSRTENRDPLPNPWA<br>PPPATQSSATTSTTTSTGSGSGNSSSNATGNTVAAANYVASIFSTPG<br>MQSLLQQITENPQLIQNMLSAPYMRSMMQSLSQNPDLAAQMMLNS<br>PLFTANPQLQEQMRPQLPAFLQQMQNPDTLSAMSNPRAMQALMQI<br>QQGLQTLATEAPGLIPSFTPGVGVGVLGTAIGPVGPVTPIGPIGPIVP<br>FTPIGPIGPIGPTGPAAPPGSTGSGGPTGPTVSSAAPSETTSPTSES<br>GPNQQFIQQMVQALAGANAPQLPNPEVRFQQQLEQLNAMGFLNR<br>EANLQALIATGGDINAAIERLLGSQPS |
| UBQLN2<br>ΔSTI1-II<br>(residues<br>379-462<br>removed) | MAENGESSGPPRPSRGPAAAQGSAAAPAEPKIIKVTVKTPKEKEEF<br>AVPENSSVQQFKEAISKRFKSQTDQLVLIFAGKILKDQDTLIQHGIHD<br>GLTVHLVIKSQNRPQGQSTQPSNAAGTNTTSASTPRSNSTPISTNS<br>NPFGLGSLGGLAGLSSLGLSSTNFSELQSQMQQQLMASPEMMIQI<br>MENPFVQSMLSNPDLMRQLIMANPQMQQLIQRNPEISHLLNNPDIM<br>RQTLEIARNPAMMQEMMRNQDLALSNLESIPGGYNALRRMYTDIQE<br>PMLNAAQEQFGGNPFASVGSSSSSGEGTQPSRTENRDPLPNPWA<br>PPPATQSSATTSTTTSTGSGSGNSSSNATGNTVAAANYVASIFSTPG<br>MQSLLQQITEQTLATEAPGLIPSFTPGVGVGVLGTAIGPVGPVTPIGP<br>IGPIVPFTPIGPIGPIGPTGPAAPPGSTGSGGPTGPTVSSAAPSETTS<br>PTSESGPNQQFIQQMVQALAGANAPQLPNPEVRFQQQLEQLNAM<br>GFLNREANLQALIATGGDINAAIERLLGSQPSLEHHHHHH |
| R-Neh2Dual-<br>sGFP | RDLELPPPYLPSQQDMDLIDILWRQDIDLGVSREVFDFSQRRKEYEL<br>EKQKKLEKERQEQLQKEQEKAFFAQLQLDEETGEFLPIQPAQHTQS<br>ETSGSMVSKGEELFTGVVPILVELDGDVNGHKFSVRGEGEGDATN<br>GKLTLKFICTTGKLPVPWPTLVTTLTYGVQCFSRYPDHMKQHDFFKS<br>AMPEGYVQERTITFKDDGTYKTRAEVKFEGDTLVNRIELKGIDFKED<br>GNILGHKLEYNFNSHNVYITADKQKNGIKANFKIRHNVEDGSVQLAD<br>HYQQNTPIGDGPVLLPDNHYLSTQSKLSKDPNEKRDHMVLLEFVTA<br>AGITHGMDELYK |
| R-Neh2Dual-<br>eGFP | RDLELPPPYLPSQQDMDLIDILWRQDIDLGVSREVFDFSQRRKEYEL<br>EKQKKLEKERQEQLQKEQEKAFFAQLQLDEETGEFLPIQPAQHTQS<br>ETSGSMVSKGEELFTGVVPILVELDGDVNGHKFSVSGEGEGDATY<br>GKLTLKFICTTGKLPVPWPTLVTTLTYGVQCFSRYPDHMKQHDFFKS<br>AMPEGYVQERTIFFKDDGNYKTRAEVKFEGDTLVNRIELKGIDFKED<br>GNILGHKLEYNYNSHNVYIMADKQKNGIKVNFKIRHNIEDGSVQLAD |

| | |
|---|---|
| | HYQQNTPIGDGPVLLPDNHYLSTQSALSKDPNEKRDHMVLLEFVTAAGITLGMDELYK |
| R-Neh2Dual-ACTR-DHFR | RDLELPPPYLPSQQDMDLIDILWRQDIDLGVSREVFDFSQRRKEYELEKQKKLEKERQEQLQKEQEKAFFAQLQLDEETGEFLPIQPAQHTQSETSGSEQVSHGTQNRPLLRNSLDDLVGPPSNLEGQSDERALLDQLHTLLSNTDATGLEEIDRALGIPELVNQGQALTGCHLEMISLIAALAVDRVIGMENAMPWNLPADLAWFRRNTLNRPVIMGRHTWESIGRPLPGRRNIILSSQPGTDDRVTWVRSVDEAIAAAGDVPEIMVIGGGRVYEQFLPRAQRLYLTHIDAEVEGDTHFPDYEPDDWESVFSEFHDADAQNSHSYSFEILERR |
| vOTU[1] | GPMDFLRSLDWTQVIAGQYVSNPRFNISDYFEIVRQPGDGNCFYHSIAELTMPNKTDHSYHYIKRLTESAARKYYQEEPEARLVGLSLEDYLKRMLSDNEWGSTLEASMLAKEMGITIIIWTVAASDEVEAGIKFGDGDVFTAVNLLHSGQTHFDALRILPQFETDTREALSLMDRVIAVDQLTS |
| AMSH* (Stam2-AMSH) [1] | GPTANPFEQDVEKATNEYNTTEDWSLIMDICDRVGSTPSGAKDCLKAIMKRVNHKVPHVALQALTLLGACVANCGKIFHLEVCSRDFATEVRSVIKNKAHPKVCEKLKSLMVEWSEEFQKDPQFSLISATIKSMKEEGVTFPSAGSQTVAAAAKNGTSLNKNKEDEDIAKAIELSLQEQKQQYTEGGSSGGSNSESIPTIDGLRHVVVPGRLCPQFLQLASANTARGVETCGILCGKLMRNEFTITHVLIPKQSAGSDYCNTENEEELFLIQDQQGLITLGWIHTHPTQTAFLSSVDLHTHCSYQMMLPESVAIVCSPKFQETGFFKLTDHGLEEISSCRQKGFHPHSKDPPLFCSCSHVTVVDRAVTITDLR |
| oTUB1* (Ube2d2-oTUB1) [1] | GPMDFLRSLDWTQVIAGQYVSNPRFNISDYFEIVRQPGDGNCFYHSIAELTMPNKTDHSYHYIKRLTESAARKYYQEEPEARLVGLSLEDYLKRMLSDNEWGSTLEASMLAKEMGITIIIWTVAASDEVEAGIKFGDGDVFTAVNLLHSGQTHFDALRILPQFETDTREALSLMDRVIAVDQLTSGGSSGGSSGGSDSEGVNCLAYDEAIMAQQDRIQQEIAVQNPLVSERLELSVLYKEYAEDDNIYQQKIKDLHKKYSYIRKTRPDGNCFYRAFGFSHLEALLDDSKELQRFKAVSAKSKEDLVSQGFTEFTIEDFHNTFMDLIEQVEKQTSVADLLASFNDQSTSDYLVVYLRLLTSGYLQRESKFFEHFIEGGRTVKEFCQQEVEPMCKESDHIHIIALAQALSVSIQVEYMDRGEGGTTNPHIFPEGSEPKVYLLYRPGHYDILYK |

[1]DUBS (vOTU, AMSH*, oTUB1*) have a residual GP sequence after cleavage with HRV 3C protease.
